## Supplement for "Human Retina-on-a-Chip: Merging Organoid and Organ-on-a-Chip Technology to Generate Complex Multi-Layer Tissue Models"

**Supplementary Figures**

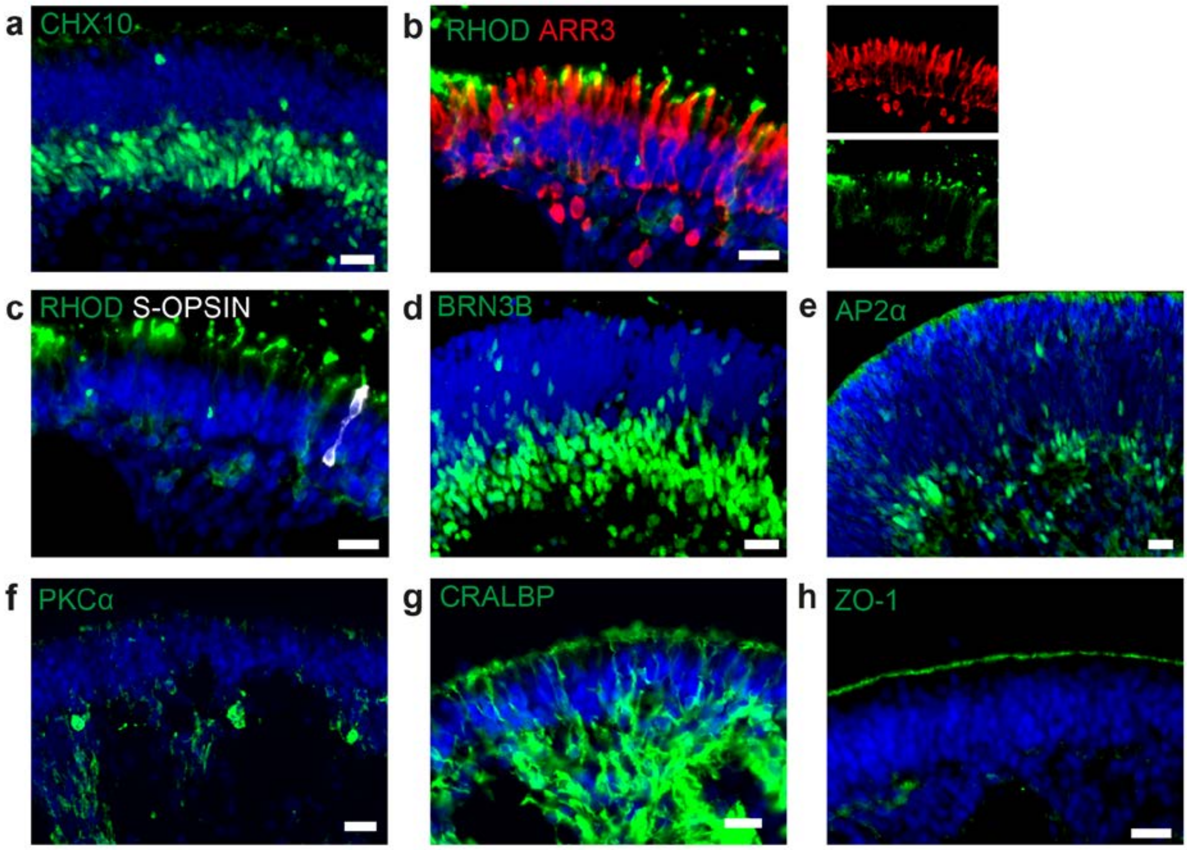


**Fig. S1 Cell types in dish cultured hiPSC-derived retinal organoids**

a) d190 RO stained for the neural retina marker CHX10 (green). b) d260 RO stained for rod marker rhodopsin (green) and cone marker arrestin 3 (ARR3, red) c) d260 RO stained for rod marker rhodopsin (green) and s-cone marker s-opsin (white). d) d42 RO stained for ganglion cell marker BRN3B (green). e) RO stained for amacrine marker AP2α (green). f) d260 RO stained for bipolar cell marker PKCα (green). g) d260 RO stained for Müller glia marker CRALBP (green) h) d260 RO stained for OLM marker ZO-1 (green). Scale bars: a-h) 20 µm. Blue: DAPI.

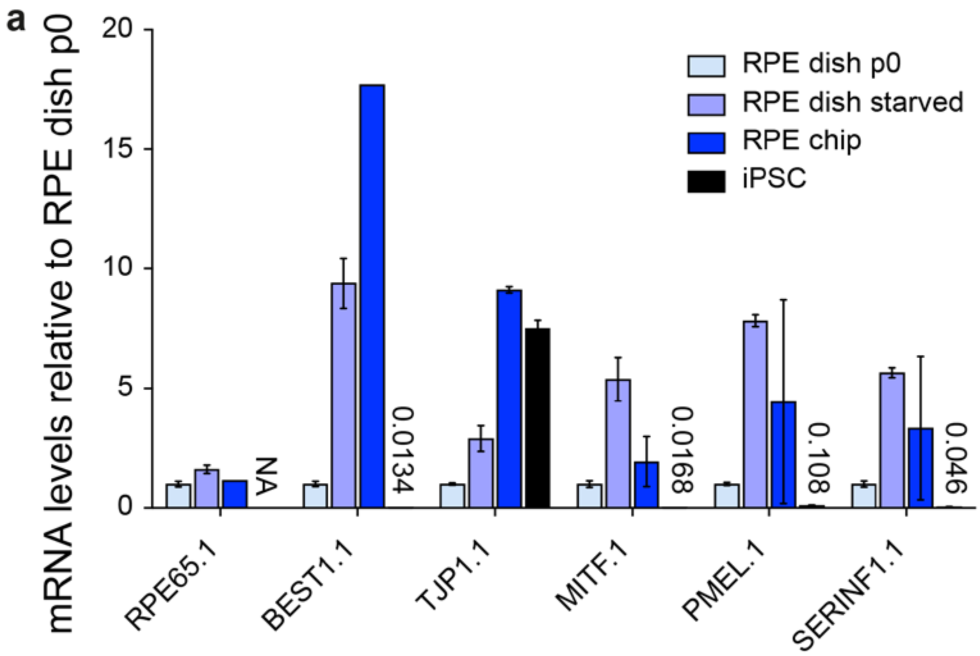


**Fig. S2 Characterization of dish and chip cultured human iPSC-derived RPE**

a) Chip cultured hiPSC-RPE immunostained for RPE markers ZO-1, RPE65 and MITF in green. b) mRNA analysis of i) dish cultured hiPSC-RPE p0, ii) after starvation for 14 days, iii) of hiPSC-RPE inside the chip and iv) respective hiPSCs. Data were normalized to dish p0 culture expression. Scale bars: a) 40 µm. Blue: DAPI. Error bars: S.E.M.

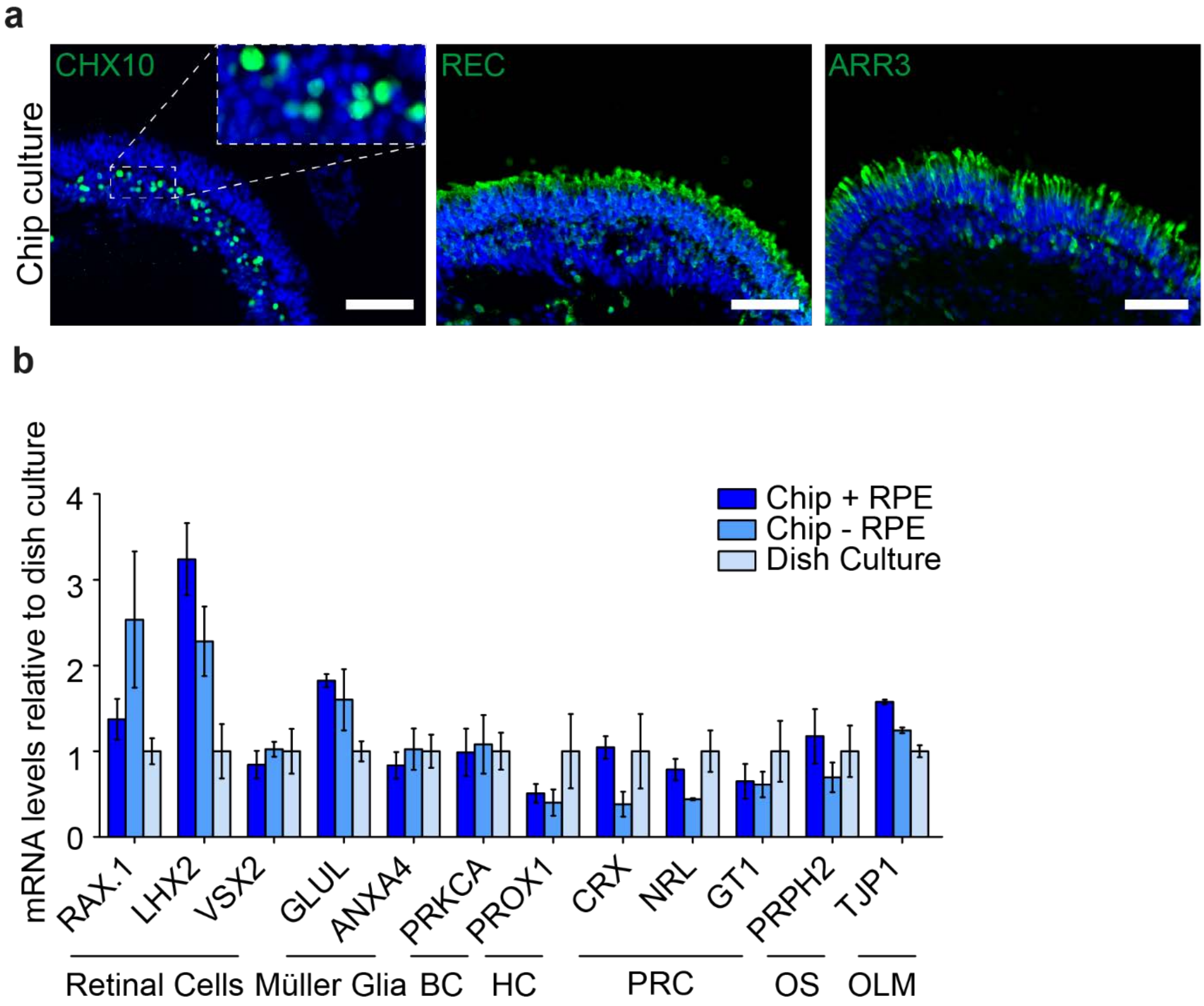


**Fig. S3 Comparison of dish and chip cultured human iPSC-derived retinal organoids**

a) After 7 days of chip-culture, d190 RO inside the retina-on-a-chip showed preserved markers for retinal cells. a1) CHX10 (green) a2) recoverin (REC, green) a3) arrestin3 (ARR3, green). a) mRNA expression from d190 organoids with and without RPE culture for 3 days inside the retina-on-a-chip were comparable to respective classically dish cultured organoids. Scale bars: a) 80 µm. Blue: DAPI. Error bars: S.E.M.

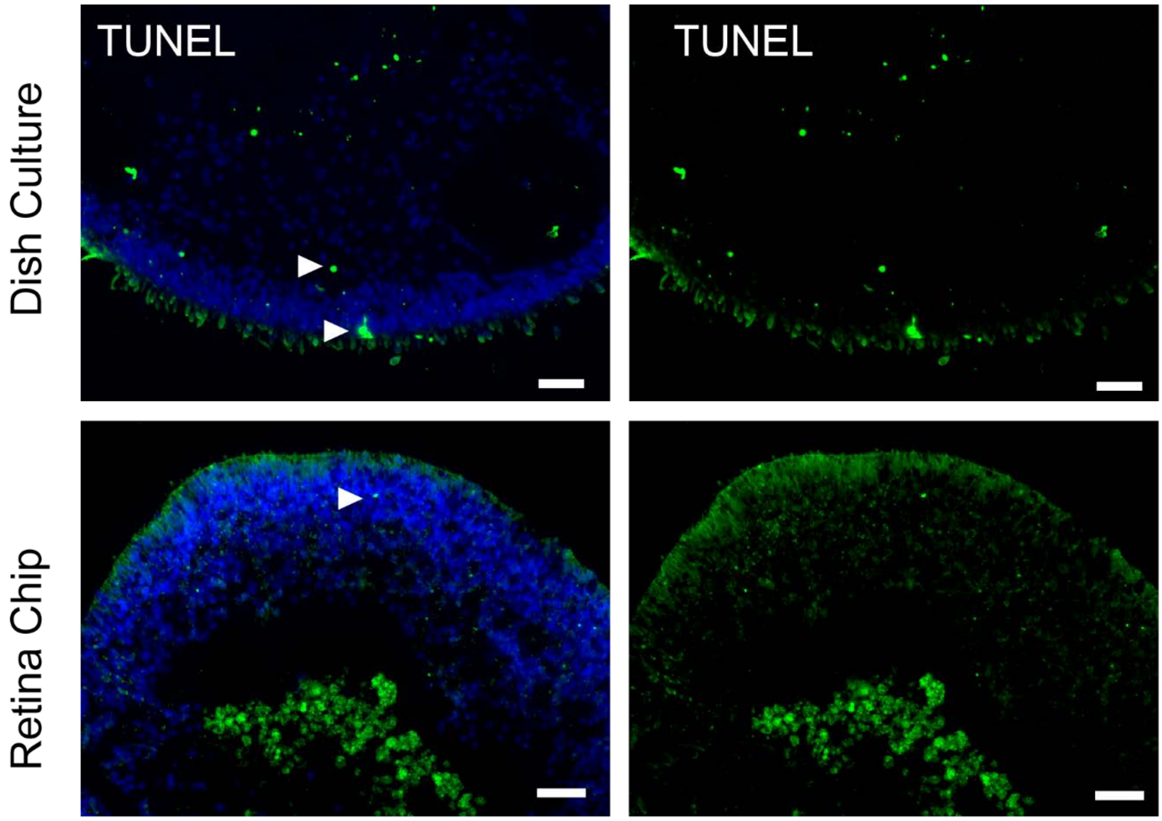


**Fig. S4 Comparison of cell death in RO cultured in the RoC or dish**
a) RO in the retina chip (upper lane) in comparison to dish culture (lower lane) labeled with the dead cell marker TUNEL. Arrows indicate exemplary positive signals. Scale bars: 40 µm. Blue: DAPI.

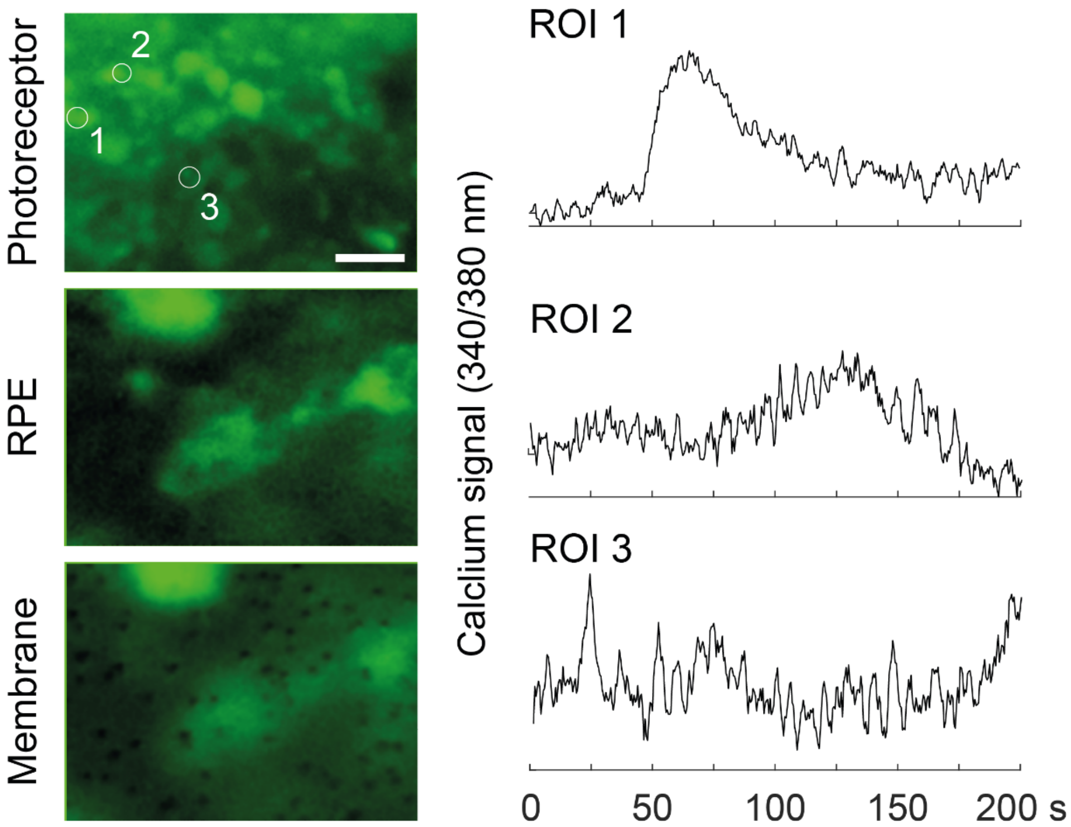


**Fig. S5 Calcium-imaging in the RoC (at 370 nm) with ratiometric calcium indicator dye Fura-2**: Outer rim of the RO at the photoreceptor layer (top), deeper focal plane at the RPE layer (middle, encircled) and the focal plane at the membrane layer, visualising the RPE contacting the membrane. Scale bar: 10 µm

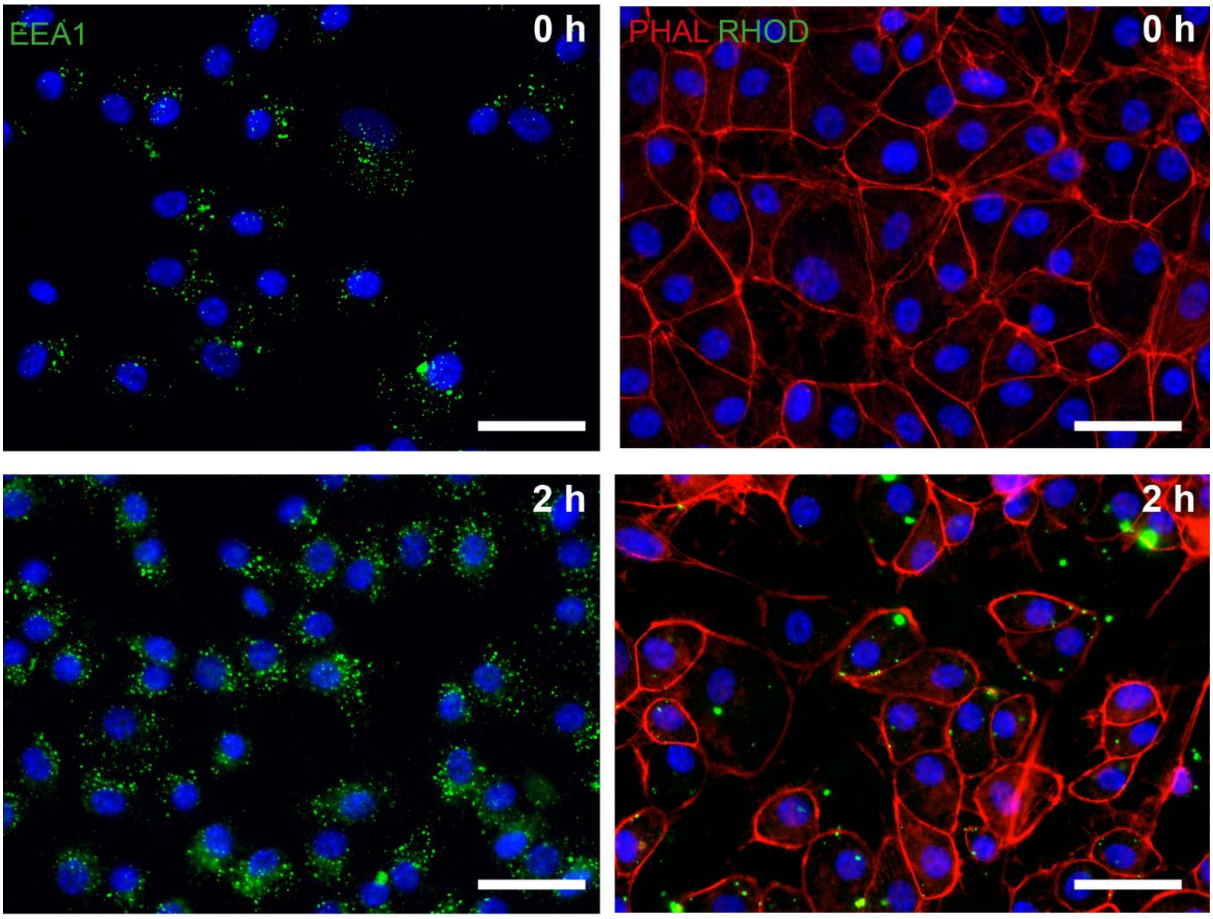


**Fig. S6 Phagocytosis assay in dish cultured hiPSC-derived RPE**

hiPSC-RPE were incubated with bovine photoreceptor outer segments (POS) and after 2 h stained positive for endosomal marker EEA1 (green, left panel) and rhodopsin (RHOD, green, right panel). Scale bars: 40 µm. Blue: DAPI.

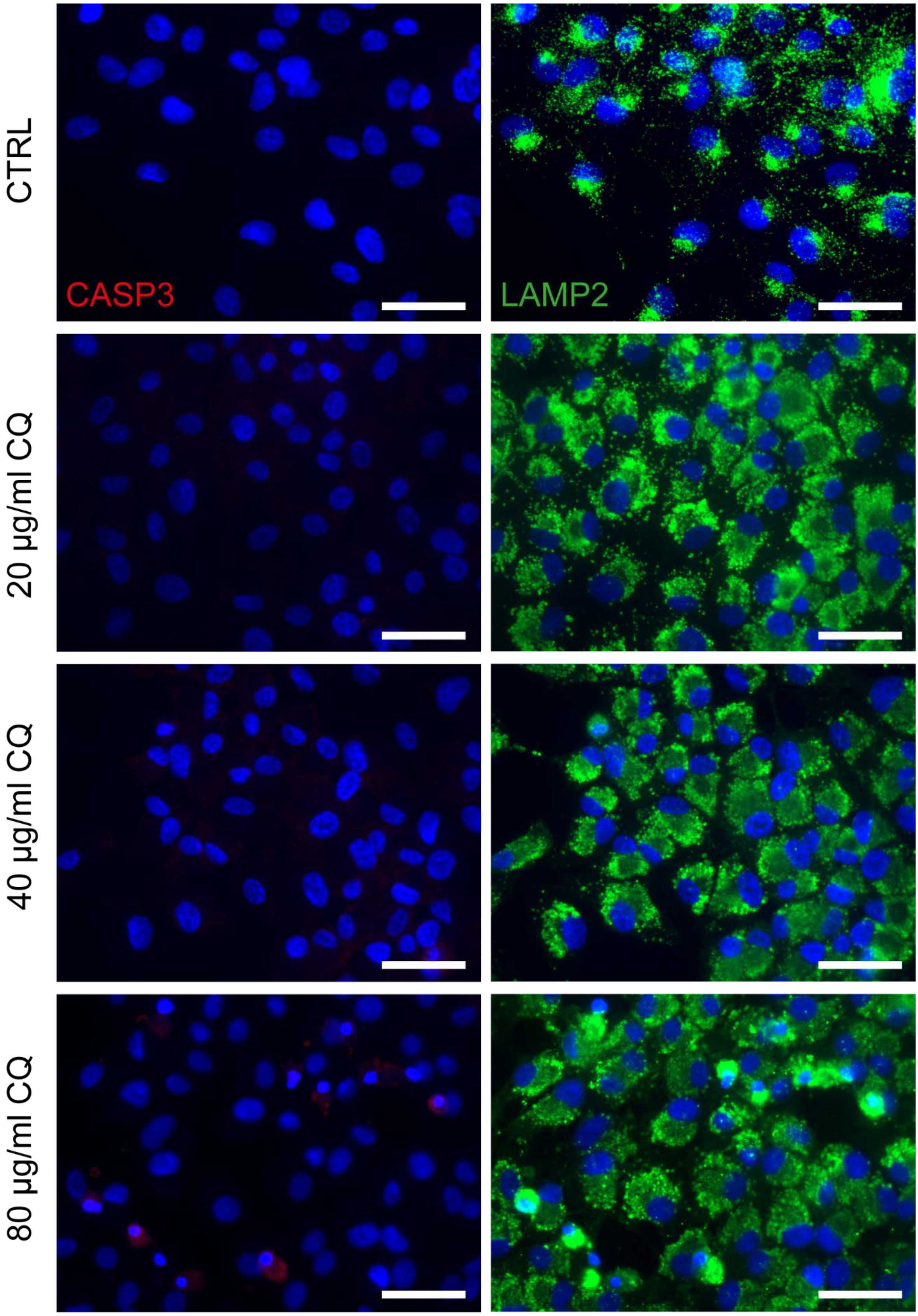


**Fig. S7 Chloroquine applied on dish culture hiPSC-RPE**
Treatment of hiPSC-derived RPE grown on cover slips with 20, 40 or 80 µg/ml chloroquine for 24 hours and immunostaining against cleaved-caspase 3 (CASP3, red) and LAMP2 (green) a) Scale bars: 40 µm. Blue: DAPI.
